# Rapid tolerance to morphine in the myenteric neurons of the small intestine is independent of β-arrestin-2 and mediated by PKC

**DOI:** 10.1101/2020.07.17.209437

**Authors:** Karan H. Muchhala, Joanna C. Jacob, Imran Alam, Shahzeb Hasan, Aliyeen Khan, Minho Kang, William L. Dewey, Hamid I. Akbarali

## Abstract

**Background and Purpose:** G-protein biased μ-opioid agonists against β-arrestin-2 activation are being investigated to reduce adverse effects. While opioid tolerance is strongly linked to the development of dependence, there is a dissociation between the two phenomena in the gut as tolerance does not develop to opioid-induced constipation, but diarrhea still manifests upon withdrawal. Here, we investigated the mechanism by which morphine tolerance in the small intestine develops.

**Experimental Approach:** Mechanism of morphine tolerance in the small intestine was evaluated *in vivo* and at the neuronal level. Whole-cell patch clamp electrophysiology was used to investigate tolerance in individual ileum myenteric neurons. Rate of morphine tolerance development in the small intestine was assessed against peripheral antinociception and whole gut transit.

**Key Results:** Tolerance develops to inhibition of small intestinal motility after one day of morphine exposure, and is more rapid compared to peripheral antinociception and constipation in chronic morphine-treated mice. Morphine tolerance was reversed by the PKC inhibitor, Tamoxifen, but not by β-arrestin-2 deletion. Similarly, β-arrestin-2 deletion did not prevent morphine tolerance to inhibition of neuronal excitability in ileum myenteric neurons. However, neuronal tolerance was attenuated by inhibiting PKC.

**Conclusions and Implications:** Unlike antinociceptive tolerance, rapid morphine tolerance in the small intestine is independent of β-arrestin-2 but is PKC-mediated. These findings reveal a potential mechanism for differences in the rates of tolerances to opioids, implicate myenteric neurons of the ileum as the primary cause for opioid-induced withdrawal effects and suggest that undesired gastrointestinal effects will persist with biased opioid agonist use.

**Summary:** **What is already known:**

- Tolerance does not develop to chronic-opioid-induced constipation but diarrhea is produced upon withdrawal
- Novel G-protein biased agonists that preclude β-arrestin-2 activation at the μ-opioid receptor are in development

**What this study adds:**

- Morphine tolerance in the ileum develops systemically and in individual myenteric neurons independent of β-arrestin-2
- Morphine tolerance in the small intestine develops before antinociception and is reversed by PKC inhibition

**Clinical significance:**

- Clinical use of G-protein biased opioid agonists will not prevent tolerance development in the ileum
- Tolerance in ileum myenteric neurons might be the basis of opioid-induced withdrawal in the gut

## Introduction

Opioid tolerance is considered as a gateway for the development of dependence (Cox, 1999). However, there is a dissociation between the two phenomena in the gut as tolerance does not develop to chronic opioid-induced constipation, but diarrhea—a common adverse effect of physical dependence—still manifests upon withdrawal (Tuteja et al., 2010; Donroe et al., 2016). We have previously shown that tolerance develops to repeated morphine exposure in the isolated ileum but not in the colon (Ross et al., 2008; Kang et al., 2012; Maguma et al., 2012) and that in myenteric plexus neurons, which regulate gastrointestinal peristalsis (Furness, 2012), naloxone induced hyperexcitability in those isolated from the ileum but not the colon of chronic morphine-treated mice (Smith et al., 2014). Given these findings, tolerance in the ileum might be associated with the induction of dependence in the gut. However, the mechanism of opioid tolerance in the ileum remains unresolved.

It is now well established that most GPCRs undergo dynamic changes in receptor confirmation to engage multiple transducers and modulatory proteins with distinct efficacies (Kenakin, 2011). This is the basis of the concept of “biased agonism” and this phenomenon has also been described for the μ-opioid receptor (MOR). MORs can engage multi-modal adaptor proteins called β-arrestins, which attenuate G-protein-mediated signaling by sterically obstructing G-protein binding to the receptor (Lohse et al., 1990; Lefkowitz, 1998). Previously, studies have shown that attenuating β-arrestin-2 (βarr2) by functional deletion or by using G-protein biased agonists results in reduced antinociceptive tolerance (Bohn et al., 2000; DeWire et al., 2013; Manglik et al., 2016; Wang et al., 2016). These studies suggest that β-arrestin-2 is necessary for MOR desensitization and ultimately opioid tolerance development.

In the gastrointestinal tract, a number of different strategies to attenuate β-arrestin-2 signaling, such as the use of β-arrestin-2 knock-out mice, phosphorylation-deficient MOR knock-in mice or G-protein biased agonists, failed to mitigate opioid-induced constipation, thus suggesting that acute opioid-induced constipation is not mediated by β-arrestin-2 (Kang et al., 2012; Maguma et al., 2012; Manglik et al., 2016; Altarifi et al., 2017; Kliewer et al., 2019). However, in the isolated colon, β-arrestin-2 deletion induced tolerance to repeated morphine, but the isolated ileum developed tolerance to morphine irrespective of the absence or presence of β-arrestin-2 (Ross et al., 2008; Kang et al., 2012; Maguma et al., 2012). Consequently, the data suggested that β-arrestin-2 mediates tolerance in the colon but its role in the development of morphine tolerance in the ileum, including in neuronal substrates that regulate motility, remains unknown. Since novel G-protein biased μ-opioid agonists are being developed for clinical use to mitigate undesired effects of opioid analgesics, it is important to delineate the mechanism of tolerance in the ileum, specifically the role of the β-arrestin-2 desensitization pathway.

The present study shows that, unlike antinociceptive tolerance, which is purportedly mediated by β-arrestin-2, morphine tolerance to small intestinal transit and in myenteric plexus neurons is mediated by a PKC-dependent mechanism that does not involve the β-arrestin-2 pathway. Furthermore, we find that morphine tolerance to small intestinal transit manifests before tolerance to antinociception and whole gastrointestinal transit. In conclusion, the present study suggests that distinct rates of morphine tolerance might be due to differences in the role of β-arrestin-2 in the mechanism of tolerance at the MOR. Furthermore, the findings presented here raise the possibility that G-protein biased agonists might induce tolerance in the ileum and lead to the development of physical dependence even if tolerance to antinociception is prevented.

## Materials and Methods

### Drugs and Chemicals

Morphine sulfate pentahydrate was obtained from the National Institutes of Health National Institute on Drug Abuse (Bethesda, MD). Pyrogen-free isotonic saline was purchased from Hospira (Lake Forest, IL). Dulbecco’s modified Eagle’s medium (DMEM)/F12, Neurobasal-A medium, Ca^2+^ and Mg^2+^ - free Hank’s balanced salt solution (HBSS), Trypsin-EDTA (0.25%), 50x B-27 supplement and L-glutamine were purchased from Gibco, Thermo Fisher Scientific (Waltham, MA). Penicillin/streptomycin/amphotericin B antibiotic-antimycotic solution and Laminin were purchased from Corning (Corning, NY). Colagenase Type 2 was purchased from Worthington Biochemical Corporation (Lakewood, NJ). Glial cell line-derived neurotrophic factor (GDNF) was purchased from Neuromics (Edina, MN). Poly-D-lysine was purchased from MiliporeSigma (Burlington, MA). Fetal Bovine serum (FBS) and bovine serum albumin (BSA) were purchased from Quality Biological, (Gaithersburg, MD) and American Bioanalytical (Canton, MA) respectively. Glass cover slips were purchased from ThermoFisher Scientific (Waltham, MA). Twenty-four-well culture dishes and 35 × 10 mm petri dishes were purchased from CELLTREAT (Pepperell, MA). CaCl_2_, MgCl_2_, NaCl, KCl, HEPES, EGTA, NaH_2_PO_4_, glucose, Na_2_ATP, NaGTP, L-aspartic acid (K salt), KOH, NaOH, MgSO_4_, NaHCO_3_, Bisindolylmaleimide (Bis) XI, charcoal, carmine red dye, tamoxifen, corn oil, and carboxymethylcellulose was purchased from MilliporeSigma (Burlington, MA). Gum arabic was purchased from Acros Organics (Fairlawn, NJ).

### Animals

All animal care and experimental procedures were conducted in accordance with procedures reviewed and approved by the Institutional Animal Care and Use Committee at Virginia Commonwealth University in compliance with the US National Research Council’s Guide for the Care and Use of Laboratory Animals, the US Public Health Service’s Policy on Humane Care and Use of Laboratory Animals, and Guide for the Care and Use of Laboratory Animals. All experiments are reported in compliance with the ARRIVE 2.0 guidelines (Lilley et al., 2020) and with the recommendations made by the *British Journal or Pharmacology*.

Swiss Webster (SW) male mice (Envigo, Indianapolis, IN), male and female β-arrestin-2 (βarr2) knock-out (KO) or their wild-type (WT) littermates, and female MOR-mCherry mice (The Jackson Laboratory, Bar Harbor, ME) were separately group-housed by genotype with up to five animals per IVC cage in animal-care quarters maintained on a 12-hour light/dark cycle. Mice used in experimental procedures were at least 7 weeks of age. Male SW mice weighed 25-35 g, male β-arrestin-2 mice weighed 20-25 g, and female β-arrestin-2 mice weighed 19-23 g. Mice were acclimated in the vivarium for at least one week prior to experimentation. Breeding pairs for the β-arrestin-2 mice were initially obtained from Dr. Lefkowitz (Duke University, Durham, NC) and housed within the transgenic facility at Virginia Commonwealth University. The genetic background of the β-arrestin-2 WT and KO mice used in our experimental procedures was 81% C57B6J:19% C57B6N. MOR-mCherry reporter mice were generously provided by Dr. J. González-Maeso (Virginia Commonwealth University). Mice had access to food and water *ad libitum* unless specified otherwise.

### *In vivo* morphine treatment

Subcutaneously implanted continuous-release morphine pellets were used to model chronic *in vivo* exposure as pellets maintain a high plasma level of morphine over a longer period of time compared to intermittent injections or osmotic pumps and the tolerance induced is more robust (Dighe et al., 2009; McLane et al., 2017). Where indicated, one 75 mg, or one or two 25 mg morphine pellets were implanted subcutaneously in the dorsum of male SW, or female or male β-arrestin-2 mice for 7 days, respectively. The rationale for using one 25 mg and two 25 mg morphine pellets in female and male β-arrestin-2 mice, respectively, is described in the ‘Pilot studies’ sub-section of ‘Materials and Methods’. Male SW mice were implanted subcutaneously with two 25 mg morphine pellets for 3 days only in the Figure 8 experiment, the rationale for which is described in the ‘Pilot studies’ sub-section of ‘Materials and Methods’. The dose of two 25 mg morphine pellets is denoted as “50 mg” throughout the present study. In all experiments, control mice were implanted with one or two placebo pellets. In order to insert the pellet, mice were first anesthetized with 2.5% isoflurane before shaving the hair from the base of the neck. Skin was disinfected with 10% povidone iodine (General Medical Corp, Walnut, CA) and alcohol. A 1 cm horizontal incision was made at the base of the neck and one or two pellets were inserted in the subcutaneous space. The surgical site was closed with Clay Adams Brand, MikRon AutoClip 9-mm wound clips (Becton Dickinson, Franklin Lakes, NJ) and cleansed with 10% povidone iodine. Use of aseptic surgical techniques minimized any potential contamination of the pellet, incision and subcutaneous space. Mice were allowed to recover in their home cages where they remained throughout the experiment.

### Isolation of myenteric neurons from ileum

Myenteric plexus neurons were isolated from the longitudinal muscle of naïve male β-arrestin-2 WT or KO mouse ileum as described by Smith et al., 2013 (Smith et al., 2013). Mice were euthanized by cervical dislocation and the ileum was immediately transferred to cold (4⁰C) Krebs solution (in mmol·L^−1^: 118 NaCl, 4.6 KCl, 25 NaHCO_3_, 1.3 NaH_2_PO_4_, 1.2 MgSO_4_, 11 glucose, 2.5 CaCl_2_) bubbled with carbogen (95% O_2_/5% CO_2_) gas. Intestinal digesta was flushed out with cold Krebs following which the intestine was transferred to a beaker containing clean ice-cold Krebs buffer. The intestine was cut into 2-4 cm segments and each segment was threaded onto a plastic rod. Longitudinal muscle to which the myenteric plexus is attached was gently teased apart with a moist cotton applicator and collected in clean carbogen-bubbled, ice-cold Krebs. Longitudinal muscle pieces were rinsed three times in 1 mL Krebs and collected by centrifugation (350 x g; 30 seconds; 4⁰C). Longitudinal muscle sheets were minced into smaller pieces with a scissor and incubated at 37⁰C for 60 minutes with 1.3mg·mL^−1^ collagenase type 2 and 0.3 mg·mL^−1^ BSA in carbogen-bubbled Krebs. Following collagenase digestion, tissue was collected by centrifugation (350 x g; 8 minutes; 4⁰C) and supernatant was discarded. Tissue was then incubated at 37⁰C for 8 minutes in 0.05% trypsin-EDTA in Ca^2+^ and Mg^2+^ -free HBSS. Digestion was halted by neutralizing trypsin with DMEM/F12 media supplemented with 10% FBS. Tissue was filtered through a sterile Nitrex mesh and cells were collected by centrifugation at 350 x g for 8 minutes at 4⁰C. Supernatant was decanted and cells were resuspended in Neurobasal-A media supplemented with 1% FBS, 1x B-27, 10 ng·mL^−1^ GDNF, 2 mM L-glutamine, and antibiotic-antimycotic solution. Isolated cells were plated on poly-D-lysine and laminin-coated glass coverslips and maintained at 37⁰C in a humidified 5% CO_2_/air-enriched incubator. 15-18 hours later, cells were used in patch clamp experiments. Where indicated, cells were cultured in media supplemented with morphine sulfate pentahydrate for 15-18 hours (overnight) prior to patch clamp electrophysiology experiments.

Cells were also isolated from male β-arrestin-2 WT or KO mice subcutaneously implanted with 50 mg morphine pellet for 7 days or male SW mice subcutaneously implanted with 75 mg morphine pellet for 7 days using the procedure described above. Isolated cells were incubated as described above in Neurobasal-A media supplemented with growth factors and antibiotic-antimycotic solution for 15-18 hours.

### Whole-cell patch clamp electrophysiology

Coverslips with adherent cells were transferred to a microscope stage plate continuously superfused with external physiological salt solution composed of 135 mM NaCl, 5.4 mM KCl, 0.33 mM NaH_2_PO_4_, 5 mM HEPES, 5 mM glucose, 2 mM CaCl_2_ and 1 mM MgCl_2_, and adjusted to a pH of 7.4 with 1 M NaOH. Patch pipettes (2-4 MΩ) made from pulled (Model P-97 Flaming-Brown Micropipette Puller, Sutter Instruments, Novato, CA) and fire-polished borosilicate glass capillaries (World Precision Instruments, Sarasota, FL) were filled with internal physiological solution and used to form a GΩ resistance seal with the cell. The internal physiological solution was composed of 100 mM L-aspartic acid (K salt), 30 mM KCl, 4.5 mM Na_2_ATP, 1 mM MgCl_2_, 10 mM HEPES, 0.1 mM EGTA and 0.5 mM NaGTP and pH adjusted to 7.2 with 3 M KOH. 100 nM Bis XI was added to the internal physiological solution in experiments designed to test the effect of PKC inhibition on cellular tolerance. Whole-cell current clamp recordings were made at room temperature using HEKA EPC 10 amplifier (HEKA, Bellmore, NY) at a 10 kHz sampling frequency and 2.9 kHz low-pass Bessel filtering. PatchMaster v2×60 (HEKA) was used for pulse generation and data acquisition. All current clamp recordings were performed five minutes after achieving whole-cell mode to allow dialysis of internal solution. Experiments were performed only on cells with healthy morphology and stable patch. Ileum myenteric plexus neurons were held at 0 pA resting current and a 10 pA pulse was applied in 5 pA steps starting from −30 pA to assess passive and active cells properties. Previously, myenteric plexus neurons with after-hyperpolarization (AHP) have been demonstrated to respond to morphine in patch clamp experiments (Smith et al., 2012, 2014). Furthermore, in the present study we observed that MOR-mCherry-positive ileum neurons that responded to morphine in patch clamp experiments exhibited AHP (Fig. S1). Therefore, only ileum myenteric plexus neurons with AHP were tested in experiments.

Effect of morphine on excitability of ileum neurons was evaluated by superfusing external physiological solution containing 3 μM morphine and measuring action potential characteristics every two minutes for up to 16 minutes. Each coverslip was discarded following morphine exposure and the same process was repeated on a freshly-mounted coverslip. Active and passive cellular properties such as threshold potential (V_thresh_), action potential spike height, resting membrane potential (V_rest_) and input resistance (R_input_) were compared before and after 3 μM morphine exposure. Action potential threshold was defined as the voltage at which the first derivative of the membrane voltage with respect to time (dV/dt) significantly deviated from zero in the course of an action potential upstroke. It was used as the primary measure of neuronal excitability in our experiments. Cellular tolerance was assessed identically in cells either incubated with 10 μM morphine overnight or isolated from mice implanted with one 75 mg or two 25 mg morphine pellets for 7 days. Values reported were not corrected for junction potentials (~12 mV).

Within the myenteric plexus, many different subtypes of neurons distinct from one another are organized into ganglia (Smith et al., 2013; Furness et al., 2014). Since each myenteric neuron is unique, our goal was to examine the effect of acute and chronic morphine exposure on excitability of individual neurons. Thus, measures from individual neurons were considered as independent values and not replicates for data analysis even though every group contained cells isolated from 5 animals, with the exception of experiments in MOR-mCherry mice, where 3 animals were used to collect data from 6 cells. ‘n’ represents the number of neurons per group and ‘N’ denotes number of animals per group.

Where indicated, individual-neuron threshold potential values at the denoted time-points were normalized to their matched baselines to minimize the natural variance of a heterogenous population of myenteric neurons (Smith et al., 2013). Percent change in threshold potential from baseline was calculated using the formula: (Test threshold potential at a given time) – (baseline threshold potential)/ (baseline threshold potential) *100. The transformation yielded baseline values fixed at ‘0’ and no standard deviation. Therefore, in accordance with the recommendations of Curtis et al. (Curtis et al., 2018), these data were not analyzed using ANOVA.

Where specified, threshold potential change from baseline values was also calculated using the formula: threshold potential change = (Test threshold potential at 16 minutes) – (baseline threshold potential).

### Behavioral testing

All testing was conducted in a temperature and light-controlled room in the light phase of the 12-hour light/dark cycle. Mice were acclimated to the testing room for at least 15-18 hours before commencing experiments to mitigate stress to the animals and eliminate confound from potential stress-induced effects on antinociception and gastrointestinal motility (Sorge et al., 2014). All animals were randomly divided into control and treatment groups. Mice were excluded from experiments if they exhibited wounds from aggressive interactions with cage mates, since injury-induced activation of the endogenous opioid system could confound nociceptive and gastrointestinal motility assays (Corder et al., 2013). Experimenters were not blinded to the identity of the groups; however, all behavioral testing was performed and repeated by multiple experimenters (K.H.M., J.C.J., I.A., S.H. and A.K) to ensure reliability of results.

### Charcoal meal transit assay

Gastrointestinal motility through the small intestine was measured using the charcoal meal transit assay, which is widely used in preclinical *in vivo* evaluation of drug safety (Harrison et al., 2004). In this assay, mice were fasted for 4-6 hours with *ad libitum* access to water on a wire mesh to prevent consumption of feces and bedding. Fasted mice were injected subcutaneously with either saline (10 μL·g^−1^ of body weight) or morphine (10 mg·kg^−1^) 30 minutes before administering a suspension of 5% aqueous charcoal in 10% gum arabic by oral gavage at dose of 10 μL·g^−1^ of body weight. 20 minutes later, mice were euthanized by cervical dislocation and the entire small intestine from the jejunum to the cecum was resected. Distance traveled by the leading edge of the charcoal meal was measured as a percentage of the total length of the small intestine and represented as % charcoal transit.

### Total Gastrointestinal Transit Assay with Carmine Dye

Total oral to anal GI transit time was measured using the carmine red dye method as previously described (Kimball et al., 2005; Li et al., 2011). A solution of carmine red, which cannot be absorbed from the lumen of the gut, consisted of 6% (w/v) carmine red dye powder in 0.5% (w/v) carboxymethylcellulose vehicle (in ddH_2_0). Each animal was orally gavaged with 150 μl of the carmine red solution through a 21-gauge round-tipped feeding needle and immediately placed in an empty cage with no bedding. Animals had access to water, but no food, during testing. The time of gavage was recorded as T_0_. Fecal pellets were monitored in 15-minute intervals for the presence of carmine red in the feces. Total transit time was determined as the interval between T_0_ and the production of the first fecal pellet fully incorporating the carmine dye. These times are reported as Total Transit Time in minutes. The studies were capped at 300 minutes. Where noted, total GI transit was also normalized within subjects as percent maximum possible effect (%MPE) calculated as: [(Test day transit time-Day 0 baseline transit time)/ (300-Day 0 baseline transit time)] x 100, adapted from Harris and Pierson, 1964 (Harris and Pierson, 1964).

### Evaluating thermal nociception

Thermal nociception was examined using the warm-water tail-withdrawal test, which represents the sensory aspects of spinally-mediated acute pain and has been classically used to test the efficacy of opioid analgesics (Mogil, 2009). In the warm-water tail-withdrawal assay, mice were gently secured in a cloth and the distal 1/3^rd^ of the tail was immersed in a water bath warmed to 56⁰C ± 0.1⁰C or 52⁰C ± 0.1⁰C. The latency to withdraw the tail from the water was recorded. A maximum cut-off of 10 seconds was set to prevent damage to the tail. Only naïve mice with control latency between 2 and 4 seconds were used in experiments. Tolerance was assessed by challenging mice with an acute subcutaneous injection of 10 mg·kg^−1^ morphine, which was the ~ED80 dose of morphine in male mice. Morphine challenge latency was compared against baseline latency. Where indicated, antinociception was quantified as %MPE, which was calculated as follows: %MPE= [(morphine challenge latency-baseline latency)/ (10-baseline latency)] x 100, adapted from Harris and Pierson, 1964 (Harris and Pierson, 1964).

### Morphine cumulative dose response curve

Baseline tail-withdrawal latencies were recorded as desetbed above for 7-day placebo or morphine-pelleted (MP) male or female β-arrestin-2 mice. Placebo-pelleted (PP) mice were injected subcutaneously every 30 minutes with 0.5, 0.5, 1, 2, 4 and 8 mg·kg^−1^ morphine to achieve final cumulative doses of 0.5, 1, 2, 4, 8 and 16 mg·kg^−1^ morphine. The final cumulative morphine doses evaluated in morphine-pelleted mice were 2, 4, 8, 16, 32, 64 and 128 mg·kg^−1^ s.c. morphine. Tail-withdrawal latencies were recorded every 30 minutes, following which mice were immediately injected with the next morphine dose. Antinociception was quantified as %MPE as described above.

### Tamoxifen reversal of tolerance

To assess the role of PKC in the mechanism of morphine tolerance, male SW mice were implanted subcutaneously with one 75 mg morphine or placebo pellet for 7 days. On the test day (Day 7), mice were fasted for 4-6 hours on a wire-mesh with *ad libitum* access to water. Subsequently, mice were injected intraperitoneally with 0.6 mg·kg^−1^ tamoxifen in corn oil or only corn oil. 30 minutes later, 10 mg·kg^−1^ morphine s.c. was injected and an additional 30 minutes later tail-withdrawal latency was re-tested to evaluate antinociception. The tamoxifen treatment protocol was adopted from Withy et al. (Withey et al., 2017). Subsequently, each mouse received an oral gavage of 10 μL·g^−1^ charcoal meal and 20 minutes later charcoal transit was assessed.

### Analysis of published single-cell RNA sequencing data of mouse ileum myenteric plexus and dorsal root ganglia neurons

Single-cell RNA sequencing data of naïve mouse ileum myenteric neurons and dorsal root ganglia (DRG) neurons published by Zeisel et al. (Zeisel et al., 2018) available at http://mousebrain.org/genesearch.html) and Usoskin et al. (Usoskin et al., 2015); available at http://linnarssonlab.org/drg/), respectively, was analyzed to visualize the expression of indicated genes in heat maps. Ileum neurons were isolated from the myenteric plexus and clustered into ENT 1-9 by Zeisel et al., and DRG neurons from L4-L6 were isolated and clustered into NP (non-peptidergic) 1-3, PEP (peptidergic) 1-2, NF (neurofilament) 1-5 and TH (tyrosine hydroxylase) by Usoskin et al. Mean expression of *Oprm1,, Scn9a, Scn10a, and Trpv1* was normalized and plotted as a heat map using Graphpad Prism.

### Data and statistical analysis

The data and statistical analysis comply with the recommendations of the *British Journal of Pharmacology* on experimental design and analysis in pharmacology (Curtis et al., 2018). Data analysis was performed in GraphPad Prism 8.0 (GraphPad Software, Inc., La Jolla, CA). Data are expressed as mean ± SEM. All experimental groups consist of N ≥ 5 and only these were subjected to statistical analysis. Experiments with N < 5 were pilot studies used to determine dosing regimens for further experiments. The statistical tests used for data analysis are indicated in the text or figure legends. The various parametric tests used were: 2-tailed paired or unpaired Student’s t-test, and two-way ordinary or repeated measures ANOVA with Bonferroni’s post-hoc test. The Dunnett’s post-test was used if comparison of multiple groups was made with a control group. For electrophysiology data normalized to matched baselines, where the control mean was ‘0’ and no SEM generated, non-parametric analysis was performed using the Friedman test followed by Dunn’s post-test (Curtis et al., 2018). Post-hoc tests were conducted only if F was significant and there was no variance inhomogeneity. Data were statistically significant when P <0.05. Sample sizes were based on our previous studies with similar experimental protocols.

### Pilot studies

#### To establish a model of *in vivo* morphine tolerance in male and female β-arrestin-2 mice

For this purpose, we initially tested morphine dose-response in both male and female βarr2 WT mice implanted with one 25 mg morphine (or placebo) pellet for 7 days in a pilot study. As shown in Figure 1A., there appeared to be a considerable rightward shift in the cumulative morphine dose-response relationship of morphine-pelleted βarr2 WT female mice compared to that of placebo-pelleted mice on Day 7. However, low animal numbers precluded statistical analysis (Fig. 1A). Alternately, ED_50_ values of placebo-pelleted and 25 mg morphine-pelleted βarr2 WT male mice, 2.18 (1.7 - 2.78) mg·kg^−1^ and 3.23 (2.26 - 4.62) mg·kg^−1^, respectively, overlapped, indicating that the potency shift was not significant (Fig. 1A). Thus, 7 days of exposure to 25 mg morphine pellet was not sufficient to produce tolerance in male mice, but seemed to induce tolerance in female mice in the warm-water tail-withdrawal assay.

**Figure 1.**
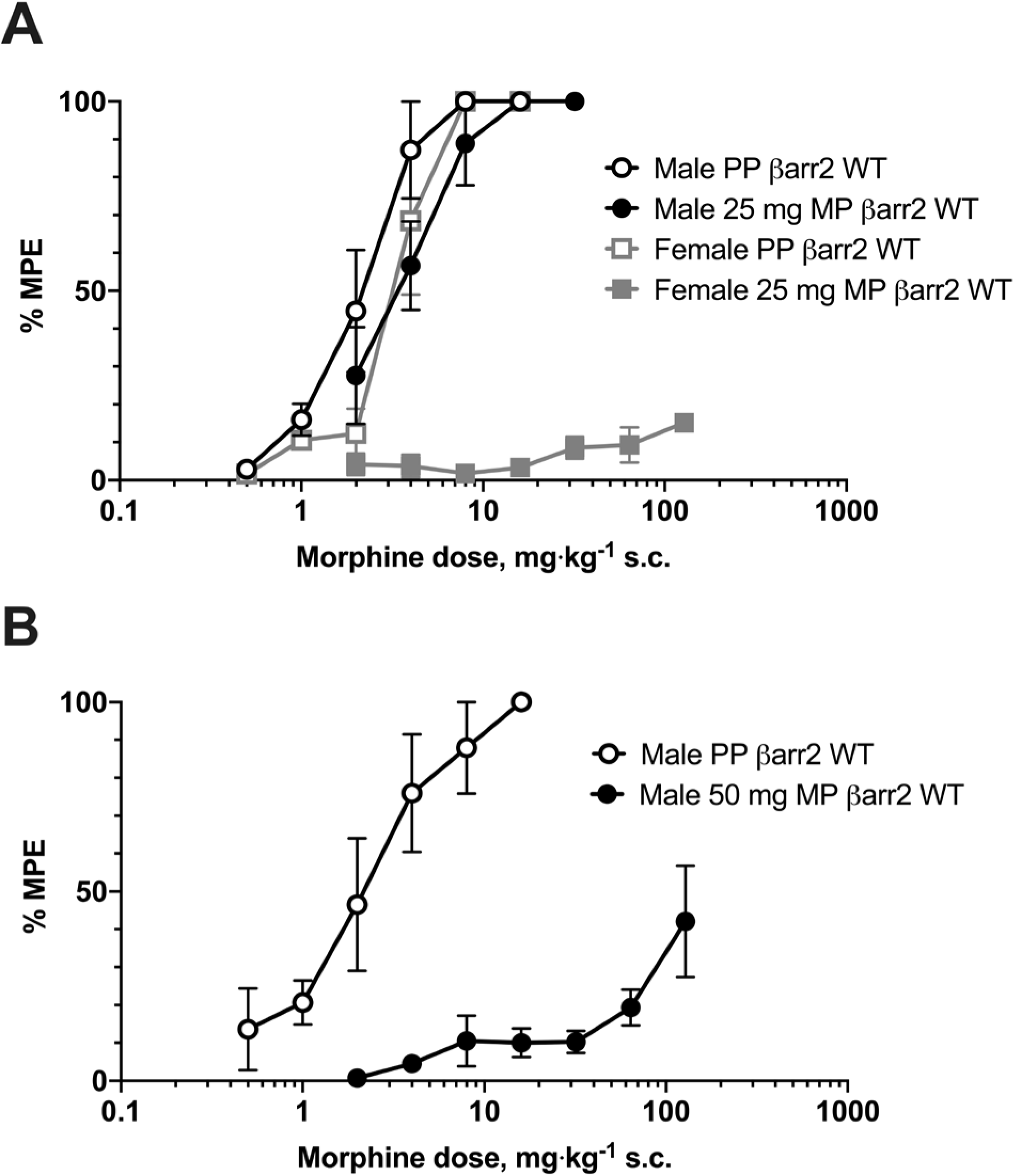
Cumulative dose response curves in morphine-pelleted male and female mice. (A) Antinociceptive tolerance represented as percent maximum possible effect (%MPE) produced by 7-day 25 mg morphine pellet (MP) in male and female mice was compared in the warm-water tail-withdrawal assay (56⁰C). 25 mg morphine pellet produced a right-shift in the morphine dose response curve of female mice, but not male mice, compared to placebo-pelleted (PP) controls. Data are mean ± SEM. Males: N=5/group for PP WT and 25 mg MP WT. Females: N=4/group for PP WT and 25 mg MP WT. (B) The morphine dose response curve of male mice was significantly right-shifted from placebo when they were pelleted with 50 mg morphine for 7 days. Data are mean ± SEM. N=6/group. *P<0.05 (versus male PP WT) by repeated measures two-way ANOVA with Bonferroni’s test.

Next, male βarr2 mice were implanted with two subcutaneous 25 mg (50 mg) morphine pellets for 7 days to test if this dose regimen induced tolerance. While the repeated measures two-way ANOVA did not reveal a significant main effect [F (3, 30) = 2.431; P = 0.085], the cumulative morphine dose response curve of male βarr2 WT mice appeared significantly right-shifted compared to placebo controls (Fig. 1B). Thus, 7-day treatment with 50 mg morphine produced tolerance to morphine-induced antinociception in the warm-water tail-withdrawal assay.

On the basis of these pilot data, subsequent experiments in βarr2 mice were carried out using 25 mg and 50 mg (2 × 25 mg) morphine pellets in female and male mice, respectively.

#### To compare the onset of morphine tolerance between antinociception and small intestinal motility

In order to evaluate the development of tolerance, it is necessary to assess whether the acute morphine challenge dose produces a response. However, we observed that in male β-arrestin-2 WT mice subcutaneously implanted with two 25 mg (50 mg) morphine pellets, the tail-withdrawal latency remained near the maximum cutoff value of 10 seconds for three days post pelleting (Fig. 2). Hence, we were unable to test tolerance to an acute morphine challenge. On the other hand, male SW mice subcutaneously implanted with two 25 mg (50 mg) morphine pellets for three days exhibited baseline tail-withdrawal latencies similar to pre-pellet, Day 0 levels (Fig. 2). Therefore, experiments to compare the rates of morphine tolerance to antinociception and in the small intestine in the same animal were carried out using male SW mice treated for 3 days with 50 mg morphine pellet.

**Figure 2.**
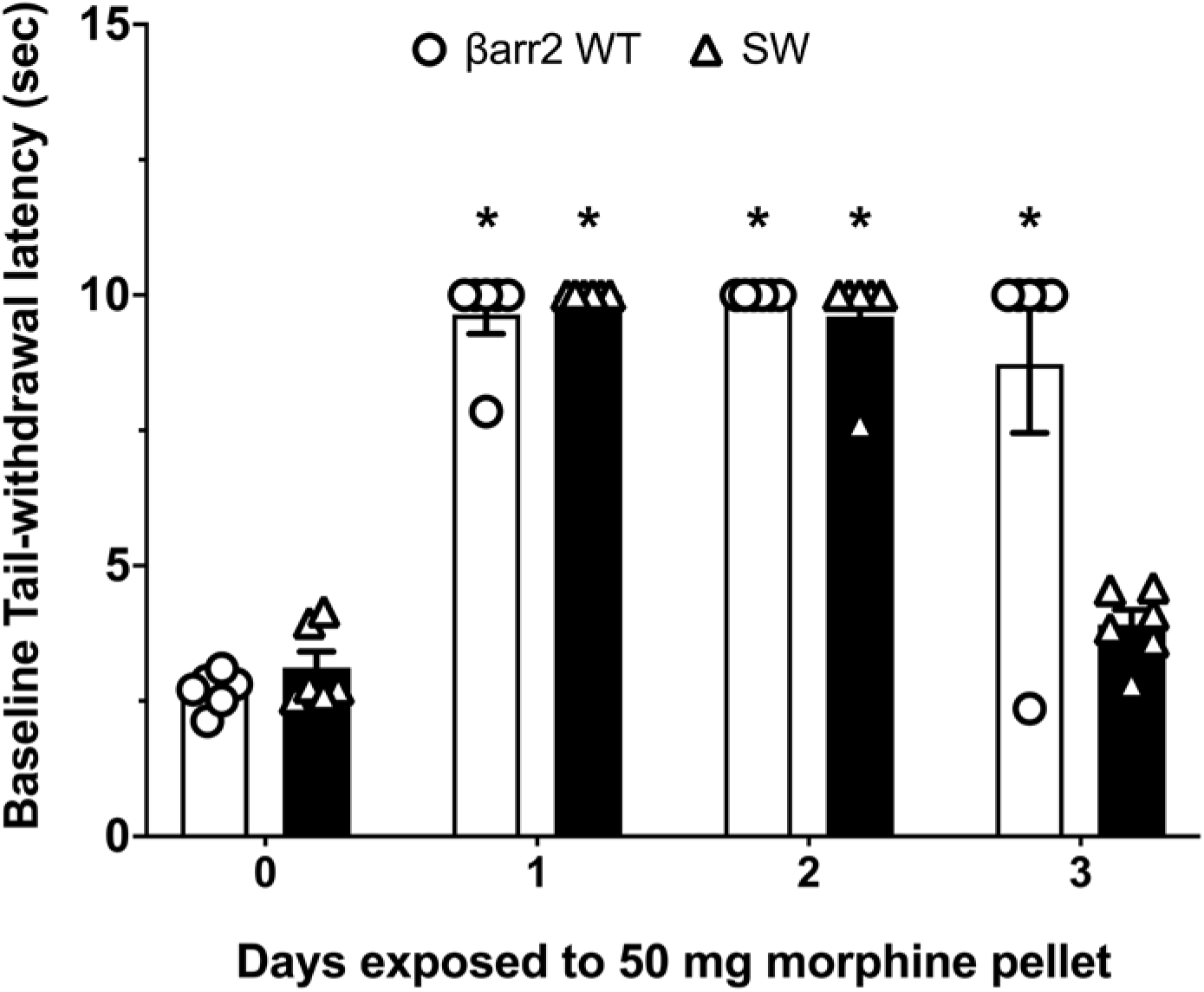
Pilot study to compare rates of morphine tolerance in small intestine and to antinociception. Daily baseline warm-water tail-withdrawal latency in male SW and β-arrestin-2 (βarr2) WT mice pelleted with two 25 (50) mg morphine pellet for 3 days. Data are mean ± SEM. Scatter represents individual animals. N=6/day. *P<0.05 versus Day 0 by two-way repeated-measures ANOVA with Dunnett’s post hoc test. Baseline latency returned to Day 0 levels in SW but not βarr2 WT mice after 3 days of morphine exposure.

## Results

### Depletion of β-arrestin-2 does not alter morphine tolerance to small intestinal motility *in vivo* in either male or female mice

We first sought to investigate whether β-arrestin-2 alters the rate at which tolerance develops to morphine-induced inhibition of motility in the small intestine *in vivo*. Furthermore, we also tested sex differences on morphine tolerance. For this purpose, we monitored the time course of the development of morphine tolerance in the small intestine of male (Fig. 3) and female (Fig. 4) βarr2 KO mice. Tolerance was induced by subcutaneously implanting male mice with two 25 mg (denoted as 50 mg) morphine pellets or female mice with one 25 mg morphine pellet as these treatment regimens produced significant morphine tolerance in the warm-water tail-withdrawal assay as reported above (Fig. 1). The development of tolerance was assessed on indicated days by challenging mice with 10 mg·kg^−1^ morphine s.c. Charcoal transit through the small intestine was used as a functional metric of motility in the small intestine. Inability of the morphine challenge dose to delay charcoal transit was described as tolerance.

**Figure 3.**
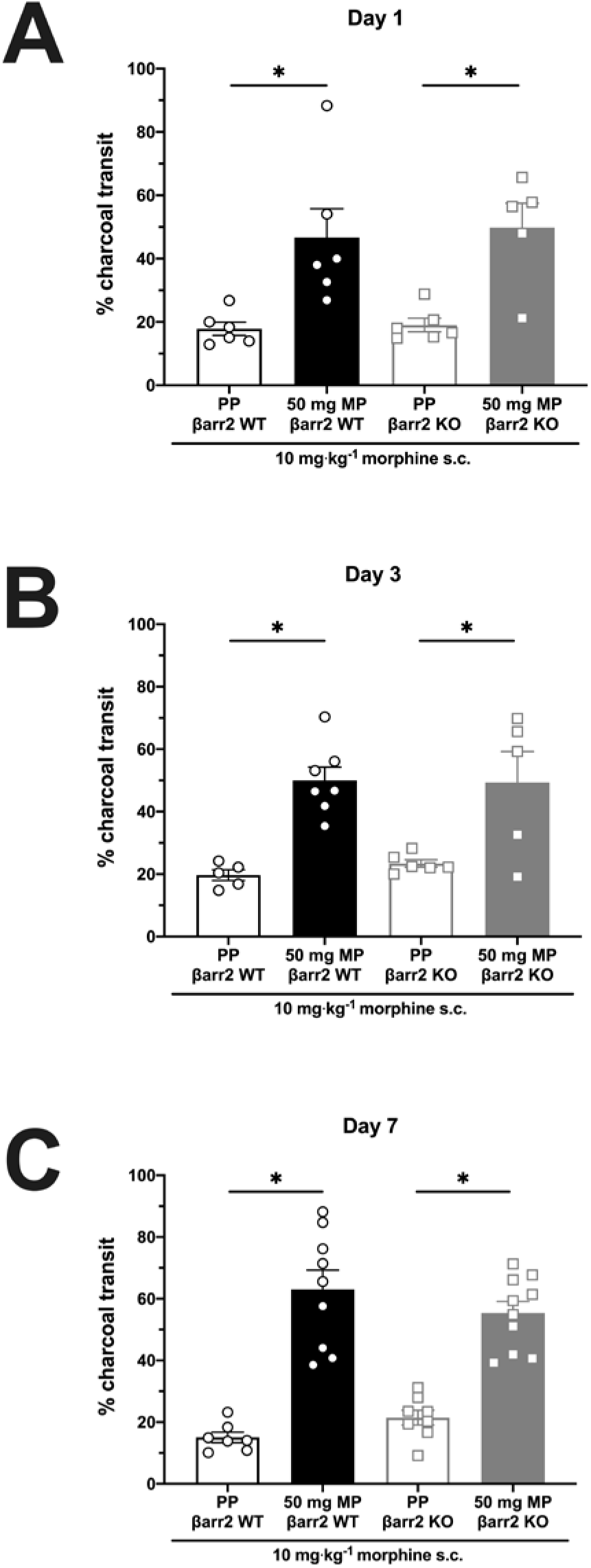
Morphine tolerance to small intestinal dysmotility in male mice is not mediated by β-arrestin-2. Morphine tolerance in the small intestine was evaluated using the charcoal transit assay in male βarr2 WT and KO mice pelleted with placebo (PP) or 50 mg (2 × 25 mg) morphine (MP) pellets. Tolerance was assessed day-wise by challenging with 10 mg·kg^−1^ morphine s.c. on indicated days. Scatter represents individual animals. Data are mean ± SEM. *P<0.05 by two-tailed unpaired t-test. In both WT and KO animals, charcoal transit was significantly greater on all test days in morphine-pelleted mice compared to placebo-controls, indicating tolerance. Ordinary two-way ANOVA did not detect a main effect of β-arrestin-2 knock out and morphine pellet treatment on charcoal transit, indicating that morphine tolerance to charcoal transit manifested irrespective of genotype.

**Figure 4.**
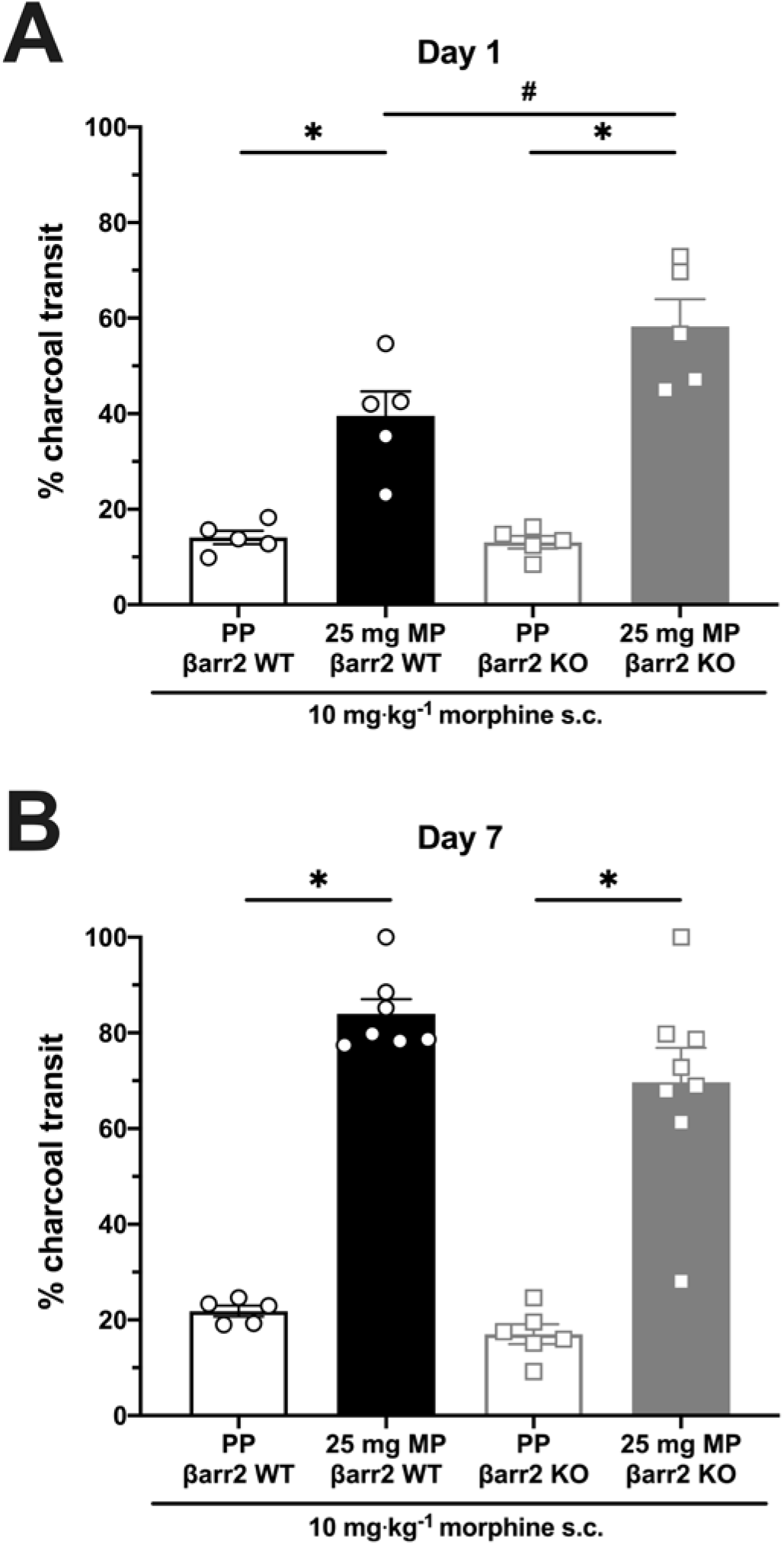
Morphine tolerance to small intestinal dysmotility in female mice is not mediated β-arrestin-2. Morphine tolerance was evaluated using the charcoal transit assay in female βarr2 WT and KO mice pelleted with placebo (PP) or 25 mg morphine (MP) pellet. Tolerance was assessed day-wise by challenging with 10 mg·kg^−1^ morphine s.c. on indicated days. Scatter represents individual animals. Data are mean ± SEM. *P<0.05 by two-tailed unpaired t-test and #P<0.05 by ordinary two-way ANOVA with Bonferroni’s post hoc test. In both WT and KO animals, charcoal transit was significantly greater on all test days in morphine-pelleted mice compared to placebo-controls, indicating tolerance. Tolerance in MP βarr2 KO mice was significantly greater than WT on Day 1, but a main effect of β-arrestin-2 knock out and morphine pellet treatment on charcoal transit was not observed on Day 7. Together, the data indicated that morphine tolerance to charcoal transit manifested irrespective of genotype.

An acute 10 mg·kg^−1^ morphine injection produced robust inhibition of small intestinal transit in male placebo-pelleted βarr2 WT and KO mice, evidenced by the reduced charcoal transit (Fig. 3). However, exposure to morphine pellet for just one day resulted in profoundly greater charcoal transit in both WT and KO mice, implicating the development of tolerance (Fig. 3A). Furthermore, two-way repeated measures ANOVA did not detect a main effect of genotype and 1 day of morphine pellet exposure on charcoal transit [F (1, 19) = 0.02683; P=0.87; Fig. 3A], suggesting that morphine tolerance was not altered by β-arrestin-2 deletion. This trend was also observed in 3- and 7-day morphine-pelleted male mice (Figs. 3B and 3C), where significantly greater charcoal transit was observed in morphine-pelleted mice compared to placebo controls post 10 mg·kg^−1^ morphine challenge, but no main effect of genotype and morphine pellet exposure on charcoal transit was found [F (1, 19) = 0.1816; P=0.67 for 3-day pelleted mice and F (1, 30) = 2.727; P=0.11 for 7-day pelleted mice].

Similar to the effects in male mice, 10 mg·kg^−1^ morphine inhibited charcoal transit in placebo-pelleted female mice with or without β-arrestin-2 (Fig. 4). However, charcoal transit was greater in 1-day morphine-pelleted female mice compared to the respective placebo controls in both genotypes, indicating the development of morphine tolerance (Fig. 4A). Surprisingly, unlike in male mice, there was a significant difference in the distance traveled by charcoal on Day 1 between morphine-pelleted WT and KO female mice (Fig. 4A). However, by Day 7 no main effect of genotype and morphine pellet exposure on charcoal transit was detected between WT and KO female mice [F (1, 22) = 0.8978; P=0.35; Fig. 4B].

In conclusion, these results suggest that β-arrestin-2 does not contribute to the development of morphine tolerance in circuits that regulate small intestinal motility of either male or female mice.

### Morphine tolerance in ileum myenteric plexus neurons is not contingent on the β-arrestin-2 pathway

In order to determine if opioid tolerance manifests in isolated neurons, whole-cell patch clamp electrophysiology in the current-clamp mode was used. For this purpose, ileum myenteric plexus neurons were harvested from β-arrestin-2 WT and KO male mice and exposed to chronic morphine *in vitro* (10 μM morphine for 15-18 hours) or *in vivo* (50 mg morphine pellet for 7 days). Cellular tolerance was assessed by monitoring changes in neuronal excitability in response to acute morphine (3 μM) challenge (Figs. 5, 6). Threshold potential, which is the membrane potential at which action potential is elicited, was used as the primary metric of neuronal excitability. Threshold potential increase from baseline signified reduced neuronal excitability. Cellular tolerance was described as the failure of the acute morphine challenge dose to alter threshold potential.

**Figure 5.**
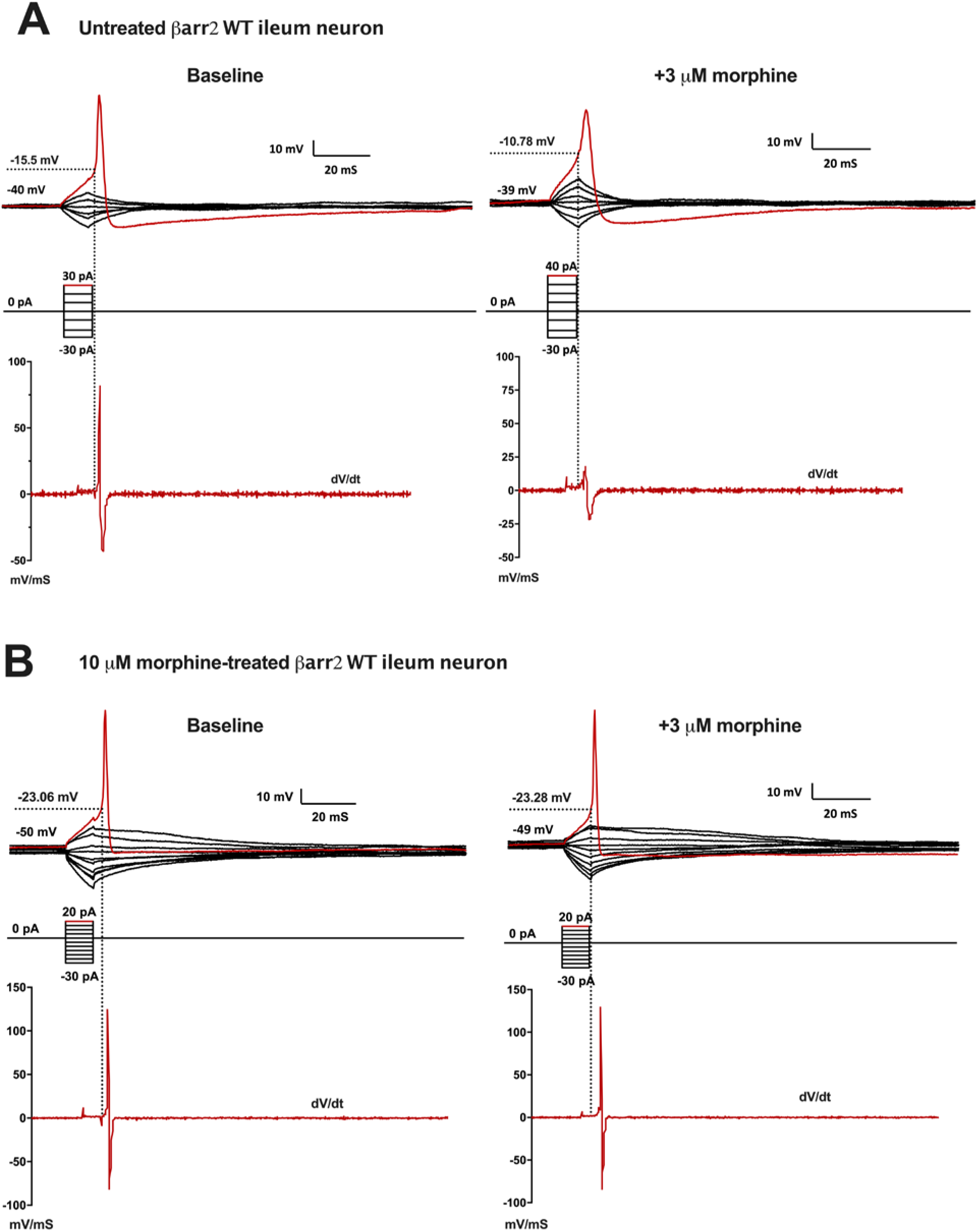
Representative current-clamp traces of untreated and 10 μM morphine-treated βarr2 WT ileum neurons. Whole-cell current clamp traces of (A) untreated or (B) 10 μM morphine-treated (*in vitr*o-treated for 15-18 hours) βarr2 WT ileum myenteric neuron at baseline (left) and after 16 minutes of acute 3 μM morphine (right). Action potential (top; in red) is generated by a step-wise current stimulation protocol (middle) with the first stimulus to elicit action potential marked in red. Threshold potential (dotted line), which is a measure of neuronal excitability, is extrapolated from the point on the action potential derivative trace (bottom), where dV/dt>0. Acute treatment with 3 μM morphine increased threshold potential of untreated βarr2 WT neuron, but not of the 10 μM morphine-treated cell.

**Figure 6.**
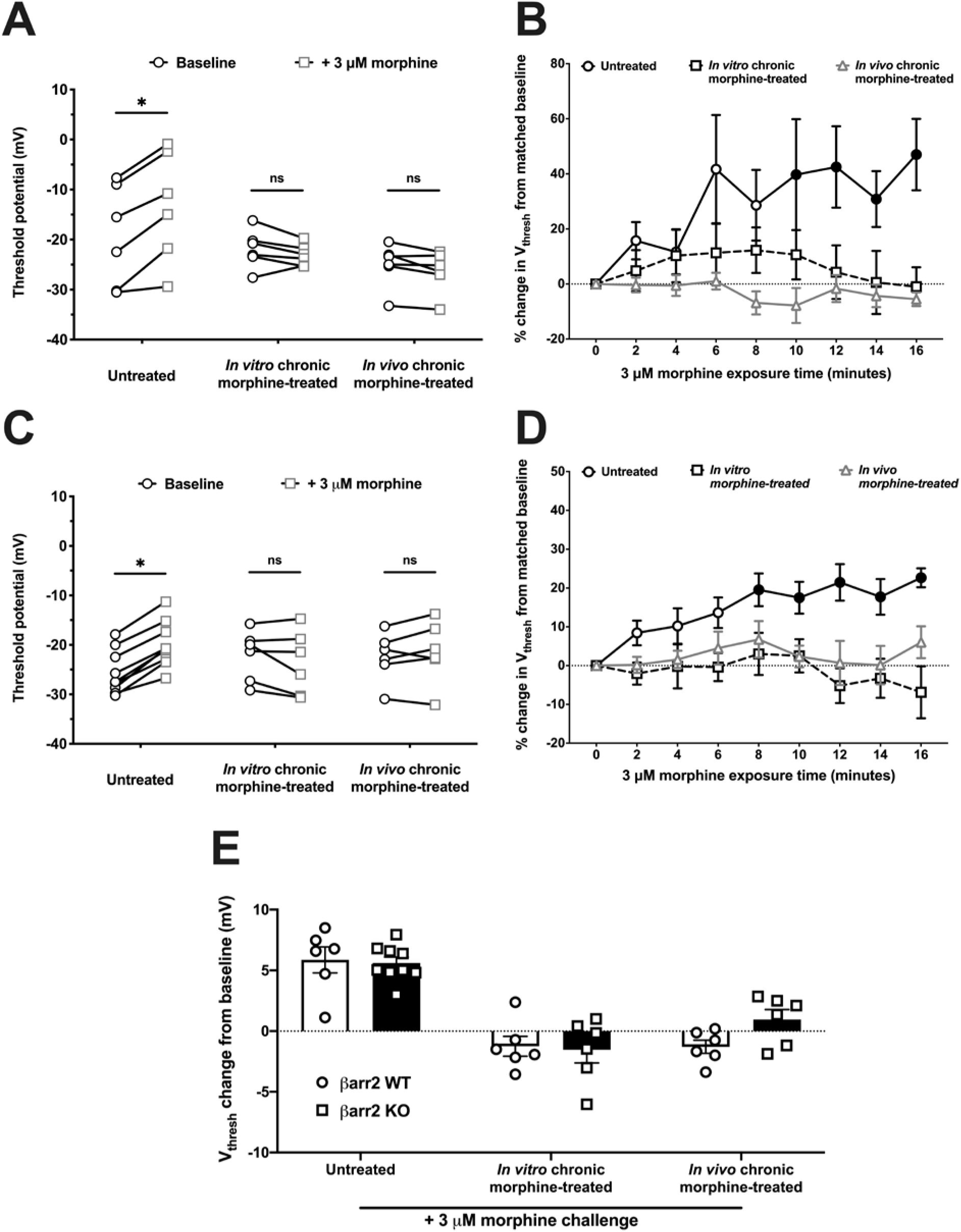
Morphine tolerance in ileum myenteric plexus neurons does not depend on β-arrestin-2. Ileum myenteric plexus neurons from male (A and B) βarr2 WT or (C and D) KO mice were bath perfused with 3 μM morphine and threshold potential (V_thresh_) changes were monitored for up to 16 minutes. (A and C) Threshold potential before (baseline) and after 3 μM morphine in individual (A) βarr2 WT or (C) βarr2 KO neurons. *P<0.05 and ‘ns’ (not significant, P>0.05) by two-way repeated-measures ANOVA with Bonferroni’s post hoc test. n= 6 βarr2 WT neurons/group; N= 5 mice/group. n=9 (untreated), 6 (*in vitro*-treated) or 6 (*in vivo*-treated) βarr2 KO neurons; 5 mice/group. (B and D) Time course of percent threshold potential change from matched baseline in (B) βarr2 WT or (D) KO neurons. n= 6 βarr2 WT neurons/group; N=5 mice/group. n=9, 5 or 6 for untreated, *in vitro* or *in vivo* morphine-treated βarr2 KO neurons; N=5 mice/group. Data are mean ± SEM and filled points represent significance from 0 minutes (P<0.05 by Friedman’s test with Dunn’s post-test). (E) V_thresh_ change across genotypes. n= 6 βarr2 WT neurons/group; N=5 mice/group, and n= 9 (untreated), 6 (*in vitro* morphine-treated) and 6 (*in vivo* morphine-treated) βarr2 KO neurons; N=5 mice/group. Data are mean ± SEM. Scatter represents individual data points. Data were analyzed with ordinary two-way ANOVA. 3 μM morphine increased threshold potential in naïve βarr2 WT and KO neurons in a time-dependent manner. The shift in threshold potential was abolished when cells were chronically treated with morphine *in vitro* or *in vivo* irrespective of genotype, indicating tolerance.

In ileum neurons from β-arrestin-2 WT and KO mice, the threshold to fire action potential significantly increased from baseline following acute exposure to 3 μM morphine (Figs. 5A, 6A and 6C). The acute morphine-induced change in threshold potential was time-dependent with significant increase from baseline observed after 10 minutes in WT mice and 8 minutes in KO mice (Figs. 6B and 6D). Notably, acute morphine exposure produced a similar change in threshold potential in naïve WT and KO neurons; ordinary two-way ANOVA failed to detect a significant main effect of genotype and pre-treatment on threshold potential change [F (2, 33) = 1.608; P = 0.22; Fig. 6E). On the other hand, no significant deviation in threshold potential from baseline was observed in *in vitro* or *in vivo* chronic morphine-treated neurons from both β-arrestin-2 WT and KO neurons following a morphine challenge (Figs. 5B, 6A and 6C). The absence of response to the acute morphine challenge persisted for 16 minutes, indicating morphine tolerance in chronically-treated cells, irrespective of genotype (Figs. 6B and 6D). Finally, acute exposure to morphine did not alter the resting membrane potential of ileum neurons (Table S1).

Thus, depletion of β-arrestin-2 does not mitigate tolerance in ileum neurons and cellular tolerance to morphine develops in ileum neurons independent of the βarr2-desensitization pathway.

### Attenuation of PKC abrogates *in vivo* morphine tolerance to small intestinal motility

It has been suggested that low-efficacy agonists such as morphine desensitize the MOR and induce tolerance through a PKC-dependent phosphorylation mechanism (Williams et al., 2013). Since morphine tolerance to small intestinal motility is not mediated by β-arrestin-2, we sought to investigate whether PKC mediates morphine tolerance to small intestinal transit *in vivo* using the charcoal transit assay (Fig. 7A). Tamoxifen was used as a PKC inhibitor in these experiments. 7-day 75 mg morphine- or placebo-pelleted male SW mice were injected with 0.6 mg·kg^−1^ tamoxifen i.p. or vehicle, following which they were challenged with 10 mg·kg^−1^ morphine s.c. to determine tolerance. 10 mg·kg^−1^ morphine reduced motility in both placebo-pelleted cohorts, as evidenced by the reduced charcoal transit (Fig. 7A). Vehicle-injected morphine-pelleted mice were resistant to 10 mg·kg^−1^ morphine-induced inhibition of charcoal transit, thus implicating tolerance (Fig. 7A). Alternately, 10 mg·kg^−1^ morphine significantly reduced charcoal transit in morphine-pelleted mice previously injected with 0.6 mg·kg^−1^ tamoxifen (Fig. 7A), indicating that morphine tolerance was reversed by tamoxifen. However, charcoal transit observed in these mice was greater than placebo-pelleted mice acutely challenged with 10 mg·kg^−1^ morphine (Fig. 7A), which suggested that the reversal of morphine tolerance in the small intestine might be incomplete.

**Figure 7.**
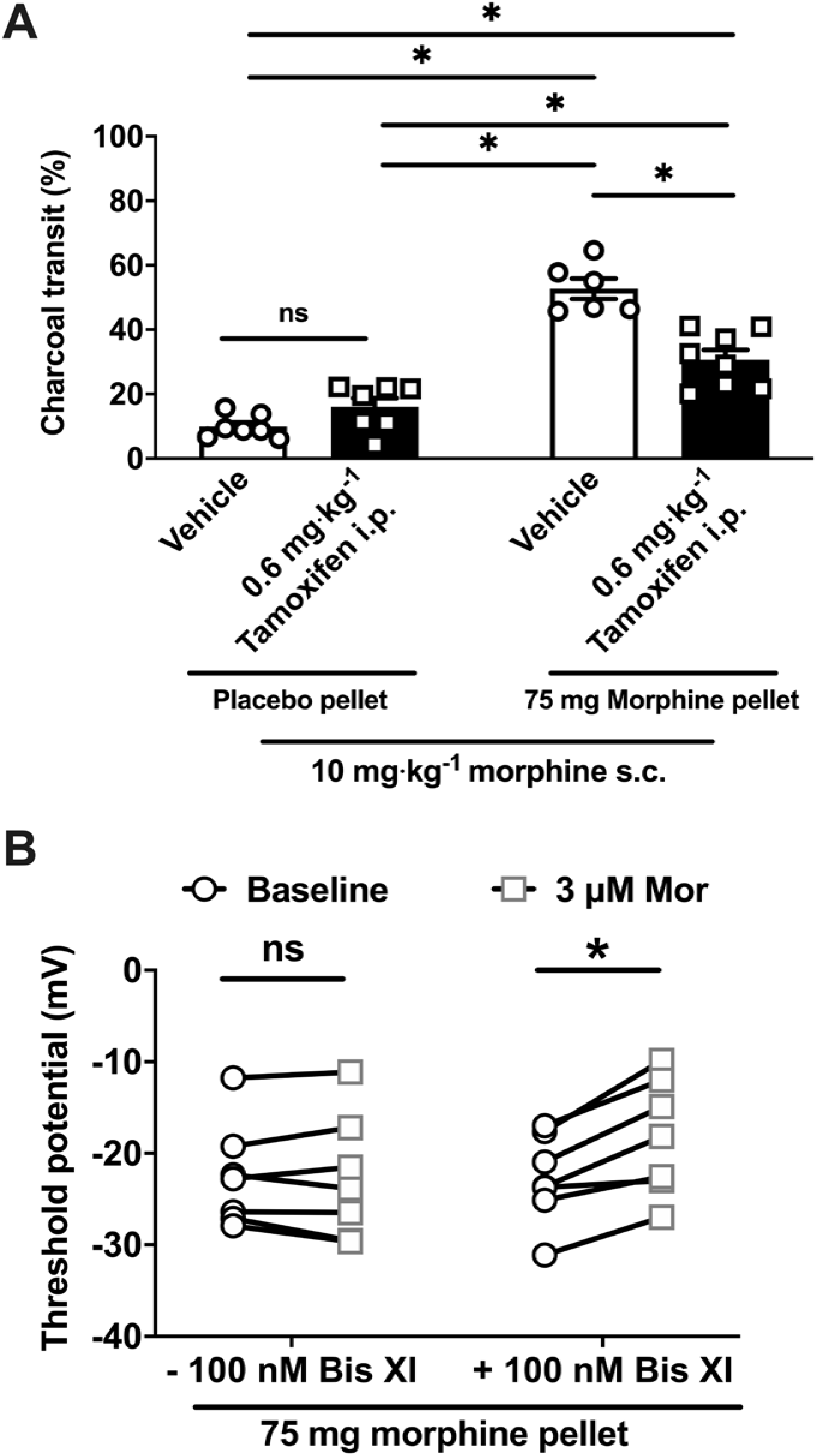
Morphine tolerance in the small intestine is reversed by acute inhibition of PKC. (A) Charcoal transit in the presence of 10 mg·kg^−1^ acute morphine challenge was measured in 7-day 75 mg morphine-pelleted (MP) or placebo-pelleted (PP) male SW mice with 0.6 mg·kg^−1^ tamoxifen or 10 μL·g^−1^vehicle i.p. Data are mean ± SEM. Scatter represents individual animals. N=7 in PP + vehicle and PP + Tamoxifen; 6 in 75 mg MP + vehicle; and 8 in 75 mg MP + Tamoxifen. *P<0.05 and ‘ns’ (not significant, P>0.05) by ordinary two-way ANOVA with Bonferroni’s post-test. 0.6 mg·kg^−1^ tamoxifen partially restored the inhibitory effect of 10 mg·kg^−1^ morphine in the small intestine of morphine-pelleted mice. (B) Threshold potential at baseline and after 3 μM morphine in individual ileum myenteric plexus neurons isolated from 7-day 75 mg MP male SW mice. *P<0.05 and ‘ns’ (not significant, P>0.05) by two-way repeated-measures ANOVA with Bonferroni’s post hoc test. N=7 cells from 5 mice/group. Neurons co-treated with 100 nM Bis XI in the pipette solution exhibited morphine-induced depolarization of threshold potential, unlike cells not exposed to Bis XI, indicating reversal of tolerance.

**Figure 8.**
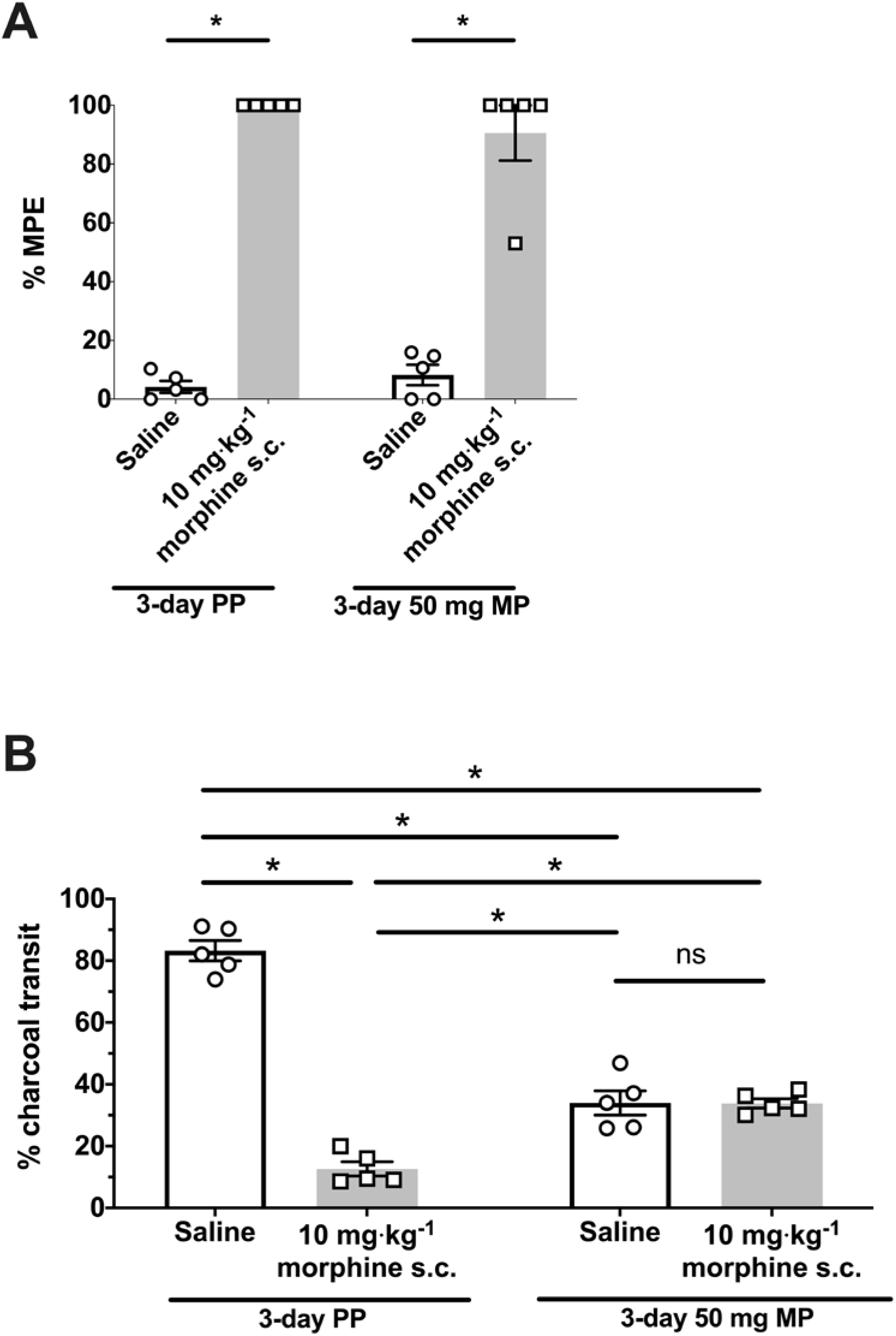
Morphine tolerance in the small intestine precedes antinociceptive tolerance. (A) Warm-water tail-withdrawal antinociception reported as percent maximum possible effect (%MPE) in 3-day 50 mg morphine-pelleted (MP) or placebo-pelleted (PP) male SW mice challenged with 10 mg·kg^−1^ morphine s.c. or 10 μL·g^−1^ saline s.c. N=5/group. Data are mean ± SEM. Scatter points are individual animals. *P<0.05 by two-tailed unpaired t-test. 10 mg·kg^−1^ morphine produced antinociception in both, placebo- and morphine-pelleted mice, compared to saline. Ordinary two-way ANOVA did not find a main effect of pellet treatment and acute morphine challenge on tail-latency, implicating the absence of antinociceptive tolerance in 3-day 50 mg MP mice. (B) In the same cohort of mice, distance traveled by charcoal through the small intestine, reported as % charcoal transit, was measured. N=5/group; *P<0.05 and ‘ns’ (not significant, P>0.05) by ordinary two-way ANOVA with Bonferroni’s post hoc test. Data are mean ± SEM. Scatter points are individual animals. The same mice exposed to morphine pellet for 3 days did not respond to the acute challenge in the charcoal transit assay, indicating tolerance to morphine-induced inhibition of small intestinal motility.

### PKC inhibition reverses morphine tolerance in the isolated ileum myenteric plexus neurons

We next investigated if morphine tolerance in the isolated ileum myenteric plexus neurons could be reversed by acute PKC inhibition (Fig. 7B). Similar to earlier electrophysiology experiments, action potential threshold was used as the primary measure of neuronal excitability, such that an increase in threshold potential signified decreased cellular excitability. Ileum myenteric plexus neurons were isolated from male SW mice pelleted with 75 mg morphine for 7 days: a paradigm that produced *in vivo* morphine tolerance in the small intestine (Fig. 7A). Cellular tolerance was determined by measuring threshold potential change in response to acute 3 μM morphine challenge. No significant shift in threshold potential from baseline in response to the morphine challenge dose was described as cellular tolerance.

In whole-cell current clamp electrophysiology experiments, threshold potential of *in vivo* chronic morphine-treated ileum neurons did not change when exposed to the acute morphine challenge (Fig. 7B), implicating cellular tolerance. No change in resting potential was detected for baseline versus 3 μM morphine challenge (Table S2). When neurons exposed to chronic morphine *in vivo* and 100 nM Bis XI, a PKC inhibitor, in the internal patch pipette solution were acutely challenged with 3 μM morphine, threshold potential depolarized and morphine tolerance was abolished (Fig. 7B). Morphine also decreased height of the action potential of Bis XI-treated cells, but not of untreated cells (Table S2). Interestingly, resting membrane potential depolarized in these cells following acute morphine exposure (Table S2). However, input resistance, which is a measure of resting ion channel activity, remained unchanged by morphine (Table S2).

These results implicate a PKC-mediated cascade, not coupled to β-arrestin-2 downstream, in the neuronal mechanism of morphine tolerance in the small intestine.

### Morphine tolerance develops to small intestinal transit more rapidly than to antinociception

We next sought to determine whether there was a difference in the time course of tolerance development between morphine-induced inhibition of small intestinal transit and antinociception. Tolerance to the inhibition of motility in the small intestine and antinociception were assessed in the same male SW mice subcutaneously implanted with two 25 mg (denoted as 50 mg) morphine pellets for 3 days using the charcoal transit assay and warm-water tail-withdrawal assay, respectively.

In the warm-water tail-withdrawal assay, an acute 10 mg·kg^−1^ morphine injection produced profound antinociception in placebo- and morphine-pelleted mice compared to those receiving 10 μL·g^−1^ saline, as evidenced by the high %MPE values (Fig. 8A). However, no main effect of acute morphine injection and pellet treatment on %MPE [F (1, 16) = 1.740; P=0.21] was detected. Consequently, this implicated that 3-day morphine-pelleted mice were not tolerant to antinociception in this assay.

In the charcoal transit assay of small intestinal motility, 10 mg·kg^−1^ morphine significantly attenuated gastrointestinal transit compared to saline controls in placebo mice (Fig. 8B). Morphine-pelleted mice challenged with saline exhibited significantly lower charcoal transit compared to placebo controls injected with saline. However, acute morphine challenge did not further diminish charcoal transit in morphine-pelleted mice, indicating that tolerance had developed to morphine-induced inhibition of small intestinal transit (Fig. 8B). Notably, morphine-pelleted mice challenged with 10 mg·kg^−1^ morphine exhibited significantly greater charcoal transit compared to placebo controls injected with morphine (Fig. 8B).

Collectively, these data show that morphine tolerance in the small intestine occurs before antinociceptive tolerance.

### Morphine tolerance to whole GI transit develops slowly compared to antinociception

Finally, we compared the rates at which antinociceptive tolerance developed versus opioid-induced constipation in male SW mice (Fig. 9). Note that the motility studies described here were capped at 300 minutes. Similar to what was previously published, we observed that the mice exhibited antinociceptive behavior in the warm-water tail-withdrawal assay on Days 1 and 2, but showed tolerance to morphine’s antinociceptive effects as early as three days following subcutaneous implantation of a 75 mg morphine pellet (Fig. 9A). In contrast, these same mice did not show any improvement in GI motility rates until Day 5, with significant constipation observed through Day 7 compared to placebo controls as assessed by the carmine dye assay (Fig. 9B). Even though GI motility rates improved significantly on Day 7 relative to day 1, there was clearly a sustained inhibitory effect of morphine that produced prolonged constipation, despite the development of tolerance to opioids and opioid effects elicited elsewhere in the body. This suggests that the prolonged constipation due to opioids might be a result of the lack of tolerance in the colon.

**Figure 9.**
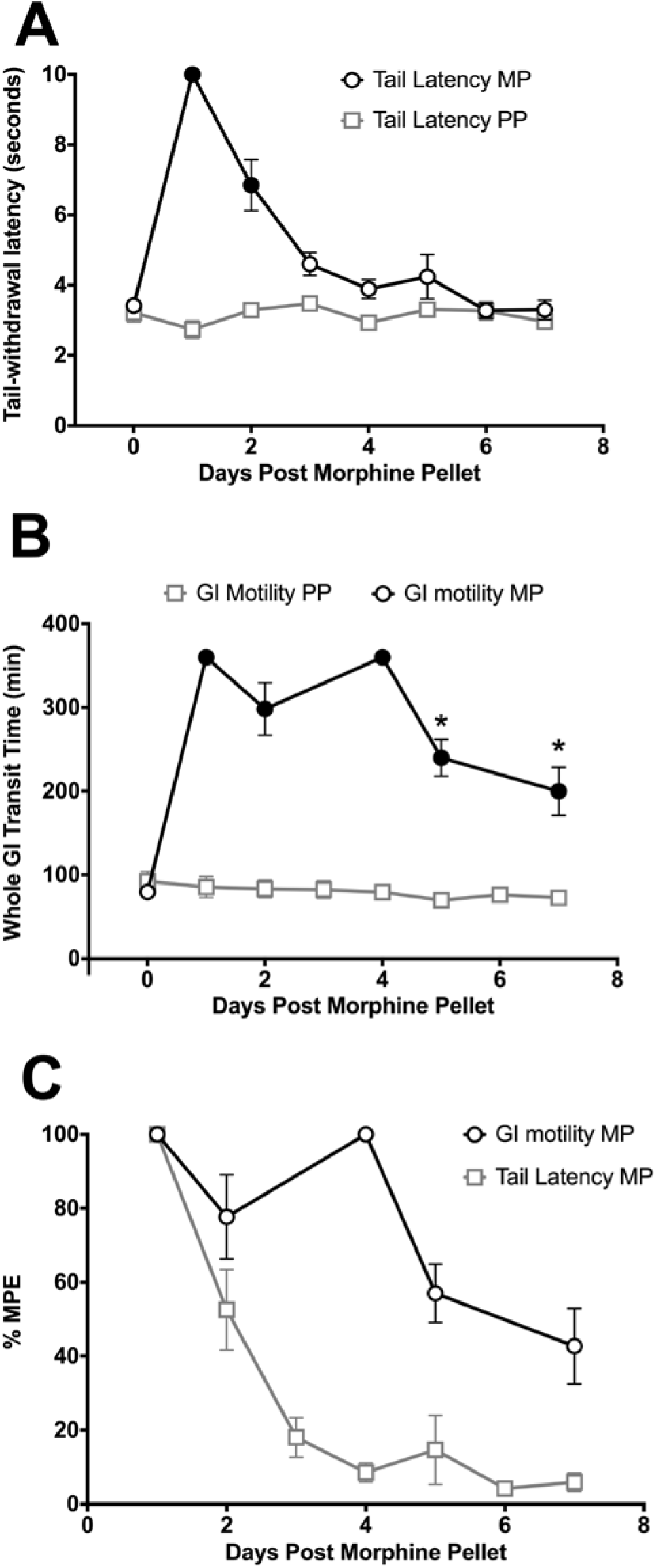
Development of morphine tolerance to antinociception and constipation. Baseline measures for tail-withdrawal latency at 52°C and total gut motility using oral carmine red dye were taken prior to implantation of a 75 mg morphine pellet (MP) (N=10) or placebo pellet (PP) (N=10). Antinociception was assessed daily using the warm-water tail-withdrawal test, beginning 24 hours post MP implantation at 52°C and ending on Day 7. In the same animals, total gut motility was assessed as the time (in min) required to pass a fresh red pellet on Days 1, 2, 3, 5 and 7. A) Antinociception was observed on Days 1 and 2 following MP implantation compared to placebo controls (Filled points are P<0.05). Tolerance to the antinociceptive effects following MP implantation developed rapidly, with no measurable morphine effect following day 3. Data are mean ± SEM, and were analyzed by two-way repeated measures ANOVA with Bonferroni’s post hoc. B) Gut motility was inhibited compared to baseline and placebo measures across all test days (Filled points are P<0.05 vs. placebo), with partial tolerance developing following day 5 (* P< 0.05 vs. Day 1 MP). Data are mean ± SEM, and analyzed by two-way repeated measures ANOVA with Bonferroni’s post hoc. C) Comparison of rate of tolerance development between GI motility and antinociception in the same animals. Data are expressed as a percent maximum possible effect (%MPE) and represent mean ± SEM, with N=10.

## Discussion

Traditional opioid agonists have a long-standing history as the most effective analgesics; however, a multitude of side effects complicate their safety margins, which become tighter as tolerance develops to the desired analgesic effect. The development of G-protein biased opioid agonists that preferentially reduce β-arrestin-2 activation have generated enthusiasm within the field and new opportunities for treatment of pain with reduced risks (Madariaga-Mazón et al., 2017; Siuda et al., 2017). While these compounds may be promising, an improvement in one opioid-dependent measurement may not equate to an overall improvement versus clinically-approved opioids.

The present study demonstrated that morphine tolerance in ileum myenteric neurons and to small intestinal transit *in vivo* develops rapidly and independently of the β-arrestin-2 pathway in both male and female mice. Moreover, morphine tolerance in neurons and to small intestinal transit is reversed by inhibiting PKC, suggesting that tolerance in the ileum is mediated by PKC. The results also show that tolerance to the same ligand—morphine—develops at a faster rate to small intestinal motility compared to spinally-mediated antinociception, tolerance to which manifests before whole GI transit. This indicates that the chronic constipation due to long-term opioid use might be a colonic effect. Taken together, the findings presented in the current study implicate that tolerance in the small intestine may develop even if β-arrestin-2 is attenuated, for example by using G-protein biased opioid agonists such as Oliceridine, which is currently awaiting FDA approval (Hertz, 2018).

Tolerance is closely associated with the development of physical dependence of which diarrhea is a common symptom (Wesson and Ling, 2003; Donroe et al., 2016). Several studies have previously reported opioid tolerance in the ileum but not in the colon, as well as precipitation of withdrawal in the isolated ileum of rodents (Collier et al., 1981; Johnson et al., 1987; Ross et al., 2008). Additionally, we have previously demonstrated that naloxone induces hyperexcitability in myenteric neurons of the ileum but not the colon of chronic morphine-treated mice (Smith et al., 2014). Furthermore, Naldemedine, a peripherally-acting MOR antagonist used clinically to mitigate chronic opioid-induced constipation, is known to commonly produce diarrhea severe enough to cause drug discontinuation (Katakami et al., 2017, 2018). Given the findings presented here that rapid tolerance to morphine-induced inhibition of peristalsis in the small intestine is β-arrestin-2-independent and in a previous study that reported intact diarrhea in morphine-withdrawn β-arrestin-2 knock-out mice (Bohn et al., 2000), it is likely that biased agonists induce dependence in the small intestine even if antinociceptive tolerance is mitigated. Consequently, the findings presented here raise questions about the usefulness of G-protein-biased agonists to mitigate adverse effects of traditional opioids.

β-arrestin-2 is a scaffolding protein that is recruited to the MOR and causes uncoupling of the receptor from G-proteins, leading to MOR desensitization (Williams et al., 2013). This desensitization pathway is widely believed to be the basis of opioid tolerance (Bohn et al., 2000). However, in the present study we observed that the rapid morphine tolerance to small intestinal transit *in vivo* is not mediated by β-arrestin-2. This finding builds upon previous studies that attenuation of β-arrestin-2 by genetic deletion or use of G-protein biased agonists, such as Oliceridine, produce tolerance to smooth muscle contractions in the isolated ileum tissue (Kang et al., 2012; Altarifi et al., 2017). Since gut motility is due to synchronous neurogenic and myogenic activity (Costa, 2000), mechanisms mediating opioid tolerance *in vivo* or *ex vivo* might not represent those underlying tolerance in the neuron. The myenteric plexus contains ganglia of neurons innervating the gastrointestinal longitudinal and circular smooth muscles that regulate peristaltic contractions (Furness et al., 2014). Here, we observed that both, “acute” and “long-term” morphine tolerance (defined by Williams *et al.*, 2013) in myenteric neurons are not mediated by β-arrestin-2. These data are consistent with *in vivo* and *ex vivo* tissue reports and indicate conclusively that opioid tolerance in the small intestine is not regulated by β-arrestin-2.

It is well-established that MOR regulation is strongly agonist dependent and it can be argued that in the present study, we used morphine, which has low affinity for β-arresin-2, to demonstrate that opioid tolerance in the small intestine is β-arrestin-2-independent. However, we have previously shown that tolerance in the isolated ileum develops to repeated exposure of high efficacy opioid agonists like fentanyl or etorphine, known to engage β-arrestins with high affinity (McPherson et al., 2010; Maguma et al., 2012; Ehrlich et al., 2019). Consequently, the β-arrestin-2-independent mechanism of tolerance in the ileum is not contingent on the opioid agonist.

Morphine, in particular, is known to preferentially induce PKC-mediated MOR phosphorylation, which has been implicated in the mechanism of *in vivo* tolerance to antinociception (Smith et al., 1999, 2006; Inoue and Ueda, 2000; Hua et al., 2002; Kliewer et al., 2019) and cellular tolerance in DRG (Jacob et al., 2018) and locus coeruleus neurons (Levitt and Williams, 2012; Arttamangkul et al., 2018). The present study showed that PKC mediates morphine tolerance to small intestinal transit *in vivo* and in myenteric plexus neurons. These findings are supported by previous observations by Wang et al., who found that PKC mediated tolerance to sufentanil-induced cAMP inhibition in chronic morphine-treated guinea pig ileum (Wang et al., 1996). Consequently, the findings presented here and elsewhere interestingly point towards a tolerance mechanism the basis of which is PKC-mediated MOR phosphorylation.

PKC is known to cause homologous or heterologous desensitization of ion channels such as voltage-gated sodium channels—pivotal to action potential generation (Levitt and Williams, 2012; Ma et al., 2019). However, we did not observe altered active action potential properties after Bis XI-induced PKC inhibition. Co-treatment with morphine was necessary for modulating neuronal excitability. This indicated that PKC-induced tolerance in ileum neurons might be due to MOR phosphorylation and not a direct effect on the ion channel. Indeed, in tolerant locus coeruleus neurons, PKC did not induce heterologous desensitization of the noradrenergic pathway (Levitt and Williams, 2012).

Numerous studies have reported contrasting sex differences in opioid-induced analgesia in both humans and rodents (Bodnar and Kest, 2010; Lee and Ho, 2013). In the present study, we found that a lesser dose of morphine induced tolerance in female mice. This could be due to the differential effect of sex hormones on the pathways underlying opioid tolerance. Indeed, a previous study reported that ovariectomized rats were refractory to morphine tolerance, indicating that estrogen might be a key regulator of opioid tolerance (Shekunova and Bespalov, 2006). While sex hormones have been noted to influence gastrointestinal physiology (Pines et al., 1998; Afonso-Pereira et al., 2018), no sex-dependent differences were noted in the mechanism of morphine tolerance to small intestinal motility.

Tolerance to opioid-induced pharmacological effects is known to develop at different rates, such that analgesic tolerance is induced faster than to respiratory depression and constipation, in humans and rodents (Ling et al., 1989; Hayhurst and Durieux, 2016; Hill et al., 2016). In this study, we use the same ligand—morphine—to demonstrate that tolerance to small intestinal transit develops before antinociceptive tolerance. Notably, in this experiment, basal small intestinal transit of morphine-pelleted animals was reduced leading to the perception of incomplete tolerance. This is likely due to opioid-induced increase in sphincter tone resulting in delayed gastric emptying (Bayguinov and Sanders, 1993; Holzer, 2009). Furthermore, morphine tolerance to whole GI transit is incomplete, implicating the colon in chronic opioid-induced constipation. Distinct rates and extent of tolerances to the same ligand raise the question of unique mechanisms of tolerance. Indeed, we find that morphine tolerance in the small intestine develops via a PKC-mediated mechanism independent of β-arrestin-2. In contrast, antinociceptive tolerance is purportedly mediated by β-arrestin-2, whereas tolerance in the colon is prevented by β-arrestin-2 (Bohn et al., 2000; Kang et al., 2012).

Voltage-gated sodium channels are critical to the action potential upstroke and altered sodium-channel kinetics in acute and chronic morphine-treated DRG and myenteric neurons have been demonstrated previously (Chen et al., 2012; Smith et al., 2012, 2014; Mischel et al., 2018). Analysis of single-cell RNA sequencing data from Zeisel at al. (Zeisel et al., 2018) and Usoskin et al. (Usoskin et al., 2015) revealed that *Scna10a* (Nav1.8 gene) is expressed only in *Trpv1* and/or *Oprm1*-containing nociceptive DRG neurons that have been previously implicated in the induction of antinociceptive tolerance (Chen et al., 2007; Corder et al., 2017), but not in ileum myenteric neurons expressing *Oprm1* (Fig. 10). Alternatively, *Scn9a* (Nav1.7 gene) is expressed in both ileum and DRG neurons. Thus, discrete mechanisms of opioid tolerances could be due to differences in MOR-ion channel coupling at the cellular level.

**Figure 10.**
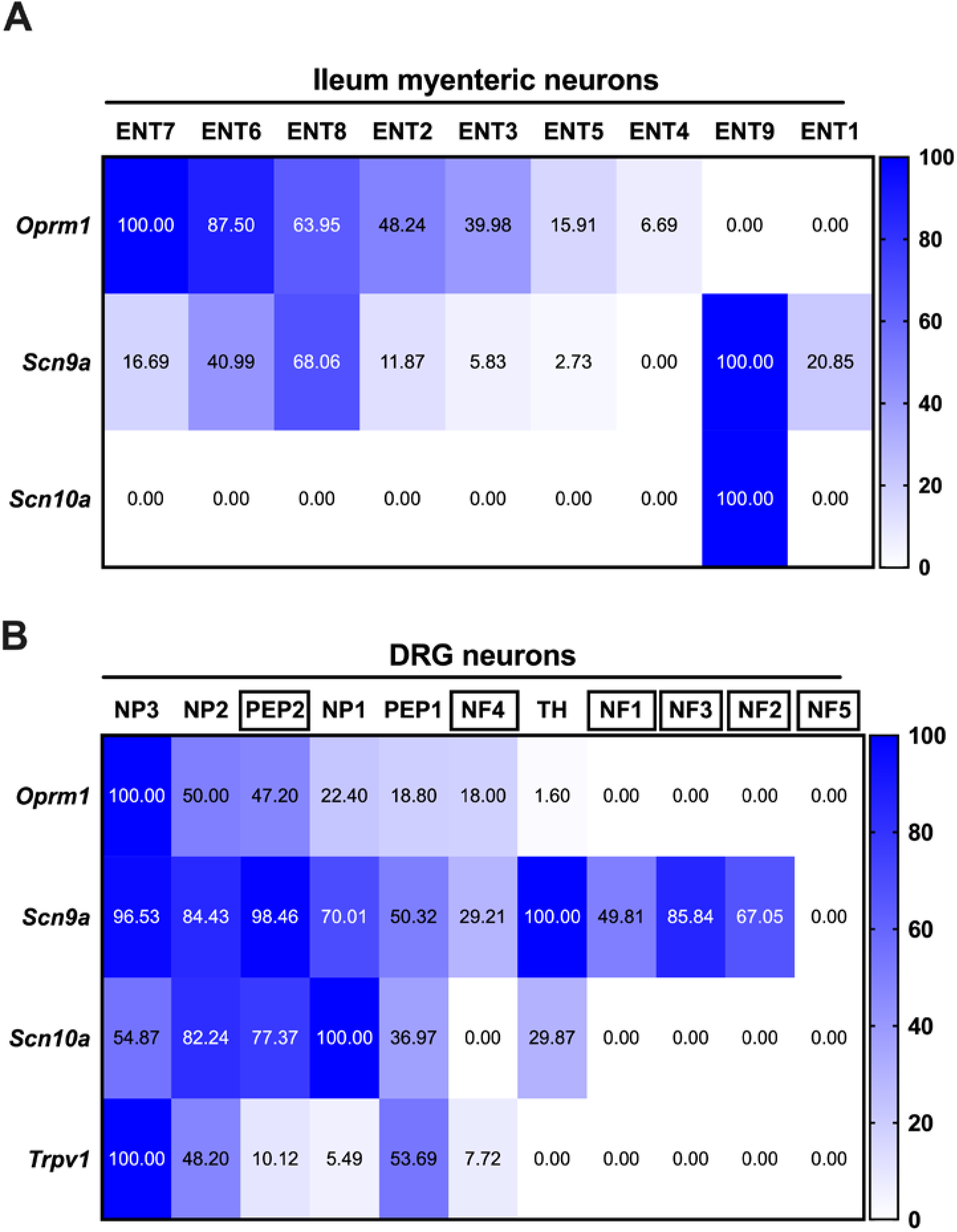
Gene expression profile of select transcripts in naïve mouse ileum and DRG neurons. Heat maps of normalized gene expression in (A) ileum myenteric plexus and (C) L4-L6 DRG neuron clusters. Correlation heat maps of genes of interest in (B) ileum myenteric and (D) DRG neurons. Single-cell RNA seq data of the mouse (A, B) ileum myenteric plexus and (C, D) DRG was obtained from Zeisel et al., 2018 and Usoskin et al., 2015, respectively. (C) ‘□’ indicates the presence of myelination in DRG neuron subsets. Gene data was normalized to the cluster expressing the most transcript.

Our conclusions from this series of studies are that in the ileum, tolerance at the MOR is extremely rapid—quicker than antinociceptive tolerance—and is mediated by a PKC-dependent mechanism not involving β-arrestin-2. This is in contrast to the finding that analgesic tolerance is caused by β-arrestin-2 (Bohn et al., 2000). This suggests that different rates of morphine tolerances could be due to discrete mechanisms in various systems. Consequently, this complicates the approach of using biased agonists as gastrointestinal dysfunction will likely remain with chronic opioid use.

## Figure Legends

**Figure. S1.**
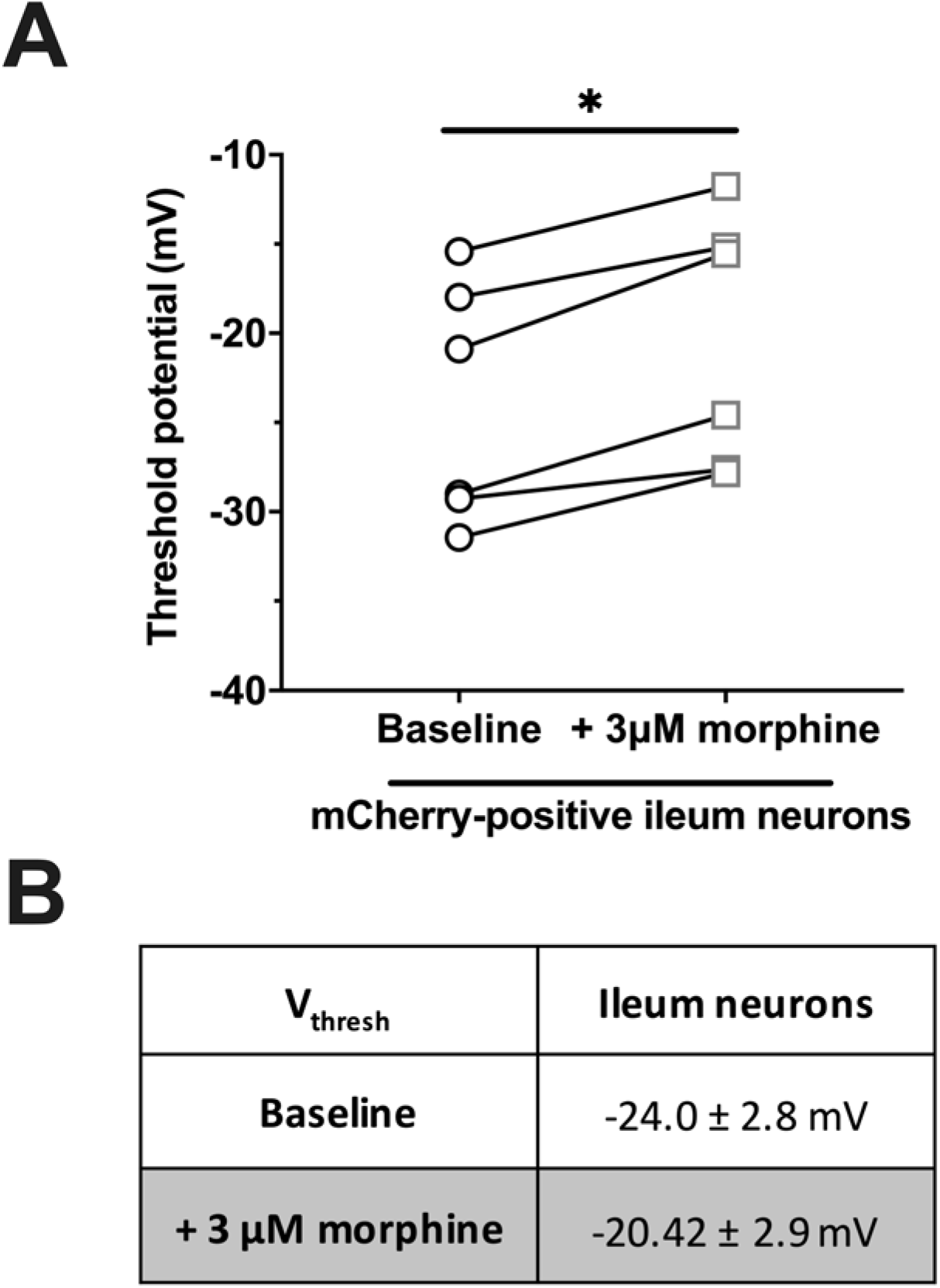
Acute morphine increased threshold potential of mCherry-positive ileum myenteric plexus neurons. (A) 3 μM morphine increased the threshold to fire action potential in mCherry-positive ileum myenteric plexus neurons. *P<0.05 by two-tailed paired t-test. All mCherry-positive ileum neurons exhibited after-hyperpolarization. (B) Action potential threshold values in mCherry-positive neurons at baseline and after 3 μM morphine exposure. Values are mean ± SEM. n=6 cells from 3 mice.

## Supplementary Table Legends

**Table S1.**
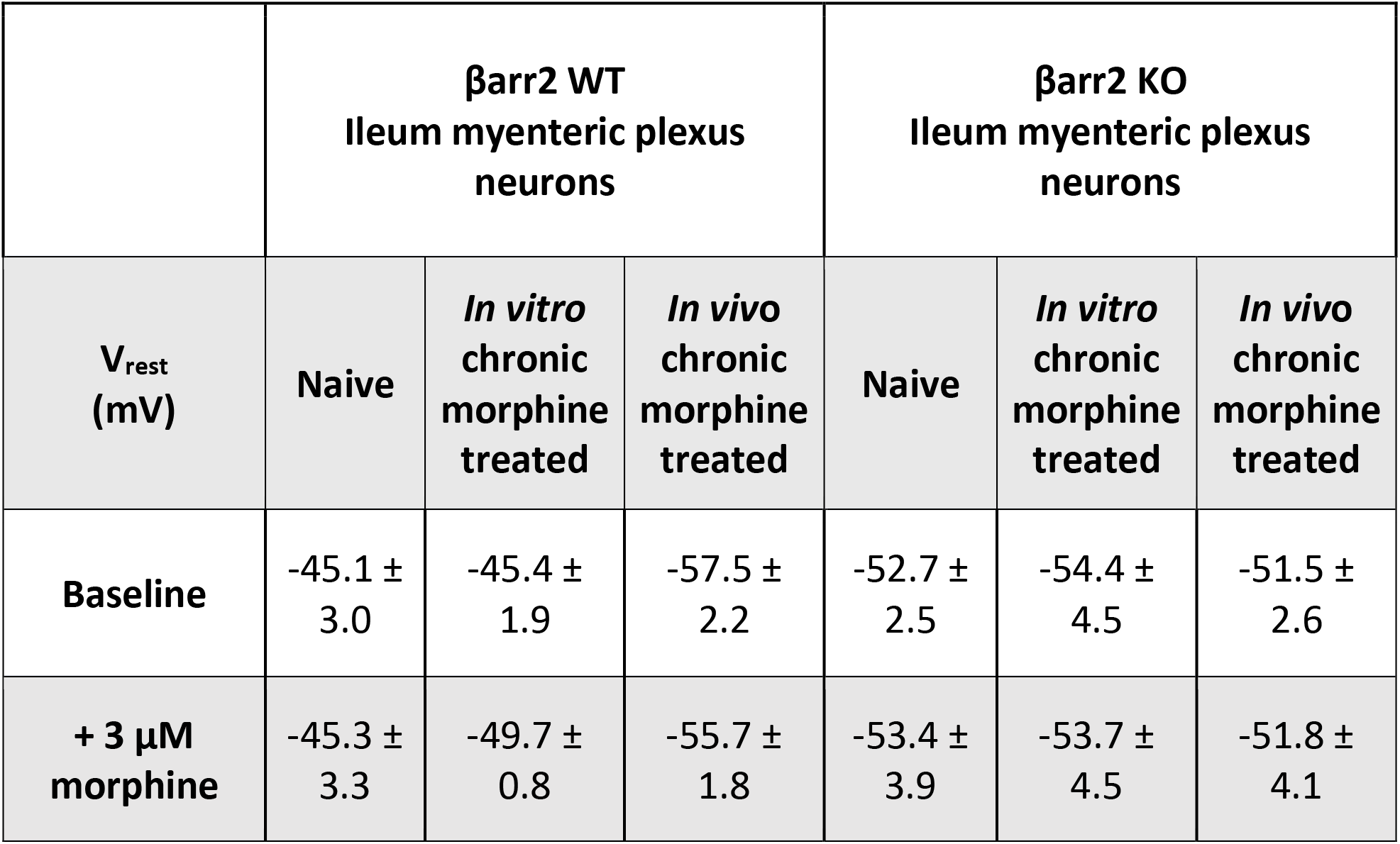
Resting membrane potential of ileum myenteric plexus neurons from male βarr2 mice. Raw resting membrane potential (V_rest_) values at baseline and after 3 μM morphine. Two-way repeated measures ANOVA did not detect a main effect of chronic morphine pre-treatment and acute morphine challenge on resting membrane potential.

| | βarr2 WT<br>Ileum myenteric plexus<br>neurons | | | βarr2 KO<br>Ileum myenteric plexus<br>neurons | | |
| --- | --- | --- | --- | --- | --- | --- |
| $V_{\text{rest}}$<br>(mV) | Naive | *In vitro*<br>chronic<br>morphine<br>treated | *In vivo*<br>chronic<br>morphine<br>treated | Naive | *In vitro*<br>chronic<br>morphine<br>treated | *In vivo*<br>chronic<br>morphine<br>treated |
| Baseline | -45.1 ±<br>3.0 | -45.4 ±<br>1.9 | -57.5 ±<br>2.2 | -52.7 ±<br>2.5 | -54.4 ±<br>4.5 | -51.5 ±<br>2.6 |
| + 3 μM<br>morphine | -45.3 ±<br>3.3 | -49.7 ±<br>0.8 | -55.7 ±<br>1.8 | -53.4 ±<br>3.9 | -53.7 ±<br>4.5 | -51.8 ±<br>4.1 |

**Table S2.** Active and passive cellular properties of ileum myenteric plexus neurons from 7-day 75 mg morphine-pelleted male SW mice. Active properties such as threshold potential (V_thresh_) and action potential (AP) height, and passive properties such as resting membrane potential (V_rest_) and input resistance (R_input_) at baseline and after 3 μM morphine. *P<0.05 vs. matched baseline by two-way repeated-measures ANOVA with Bonferroni’s post-test.

|  |  | **- 100 nM Bis XI** | **+ 100 nM Bis XI** |
| --- | --- | --- | --- |
| **V<sub>rest</sub> (mV)** | **Baseline** | -51.5 ± 2.8 | -52.1 ± 3.2 |
|  | **+ 3 μM morphine** | -51.2 ± 2.8 | -47.5 ± 2.8* |
| **R<sub>input</sub> (MΩ)** | **Baseline** | 621.1 ± 60.0 | 428.4 ± 74.3 |
|  | **+ 3 μM morphine** | 582.1 ± 61.6 | 477.0 ± 93.1 |
| **AP height from V<sub>rest</sub> (mV)** | **Baseline** | 98.8 ± 5.7 | 102.6 ± 5.5 |
|  | **+ 3 μM morphine** | 95.9 ± 7.0 | 88.1 ± 9.3* |

## Acknowledgements

The authors wish to thank David Stevens and Dr. Krista Scoggins for their technical assistance. This study was supported by the National Institute of Health grants: P30 DA033934, R01 DA036975, R01 DA024009

## Notes

**Conflict of Interest:** The authors declare no conflict of interest or competing financial interests.

### Competing Interest Statement

The authors have declared no competing interest.

